# Parental dietary vitamin B12 causes intergenerational growth acceleration and protects offspring from pathogenic microsporidia and bacteria

**DOI:** 10.1101/2023.09.05.556441

**Authors:** Alexandra R. Willis, Winnie Zhao, Ronesh Sukhdeo, Nicholas O. Burton, Aaron W. Reinke

## Abstract

The parental environment of *C. elegans* can have lasting effects on progeny development and immunity. Vitamin B12 exposure in *C. elegans* has been shown to accelerate development and to reduce killing caused by pathogenic *Pseudomonas aeruginosa*. Vitamin B12 is also maternally transferred to offspring, suggesting a potential role in progeny development. However, the intergenerational effects of vitamin B12 in *C. elegans* remain uncharacterized. Here, we show that parental exposure to dietary vitamin B12 or vitamin B12-producing bacteria results in offspring with accelerated growth. We find that this inherited acceleration phenotype is vitamin B12 dose-dependent and persists for a single generation. Progeny from vitamin B12 treated parents display reduced levels of the acdh-1 dietary sensor in the earliest larval stages, with expression of this protein recovering into adulthood. During infection with *Nematocida parisii*, a natural microsporidian pathogen, the offspring of worms fed vitamin B12 diets have better reproductive fitness. However, vitamin B12 diet does not affect offspring infection levels, suggesting that accelerated development provides tolerance to microsporidian infection. Offspring from parents exposed to vitamin B12 are also protected from killing by pathogenic *Pseudomonas vranovensis*. Vitamin B12-induced intergenerational growth acceleration and *N. parisii* tolerance is dependent upon the methionine biosynthesis pathway. However, protection from *P. vranovensis* killing is mediated through both the methionine biosynthesis and the propionyl-CoA breakdown pathways. Our results show how parental microbial diet impacts progeny development through the transfer of vitamin B12 which results in accelerated growth and pathogen tolerance.

## Introduction

Parental environment has a major impact on offspring fitness, with these effects lasting from one to many generations [1]. One common multigenerational effect is that of pathogen exposure, which has been demonstrated to increase immunity or resistance to pathogens in dozens of animals [2]. The nematode *Caenorhabditis elegans* has become an important model for understanding immune memory across generations and infection of parents by either bacteria or microsporidia, a eukaryotic parasite, primes offspring to have increased resistance to subsequent exposure [3–5]. Parental nutrition has also been shown to have a large impact on offspring immunity in many animals, but the mechanisms of this are poorly understood [6]. The interplay between immune priming, diet, and pathogen infection are largely unknown.

Vitamin B12 is an important nutrient for *C. elegans* growth and development [7]. This vitamin is missing from the standard lab diet *Escherichia coli* OP50, but when worms are grown on bacteria that produce vitamin B12 or given exogenous vitamin B12, the worms undergo accelerated development [8,9]. Many of the bacterial species that *C. elegans* encounters in nature are missing the pathways for vitamin B12 biosynthesis, but many *Pseudomonas* bacteria commonly produces this metabolite [10]. Vitamin B12 is maternally transferred to offspring through the transporter MRP-5 and worms grown without vitamin B12 for serval generations have defects in fertility and lifespan [11,12].

To understand the intergenerational impact of diet, we studied how *C. elegans* parents exposed to vitamin B12 produce offspring that are protected against pathogens. We show that offspring from parents fed vitamin B12 or bacteria that produce vitamin B12 experience accelerated growth that lasts a single generation. We demonstrate that this intergenerational accelerated growth provides tolerance to infection by the microsporidia *Nematocida parisii* and protection against killing by the pathogenic bacteria *Pseudomonas vranovensis*. There are two vitamin B12 dependant pathways in *C. elegans* and both are required for intergenerational protection to *P. vranovensis*, but only methionine biosynthesis is required for intergenerational growth acceleration and tolerance to *N. parisii*. Together our study demonstrates that bacteria can have both immune and dietary effects that are transmitted across generation which can influence infection and fitness outcomes of progeny.

## Results

### *Pseudomonas vranovensis* and dietary vitamin B12 cause intergenerational growth acceleration

Worms exposed to *P. vranovensis* produce offspring that are resistant to *P. vranovensis*-mediated killing [4]. While investigating whether parents grown on *P. vranovensis* produce offspring that might be resistant to the microsporidian parasite *N. parisii* [13], we noticed that these offspring developed more quickly than control worms. To characterize this difference in growth rate, we grew synchronized populations of *C. elegans* on non-pathogenic *Escherichia coli* strain OP50. At the L4 stage (48 h post L1), the population was split, and half of the worms replated on OP50 (as a control) and half plated on *P. vranovensis*. After a further 24 h, parent populations were treated with sodium hypochlorite to release the naïve (grown on OP50) and *P. vranovensis*-primed F1 embryos. Following hatching, synchronized F1 worms were plated on OP50 at the L1 stage. Animals were fixed at a 51 h and stained with the chitin-binding dye Direct Yellow 96 (DY96) to visualize worm embryos. Imaging by fluorescence microscopy revealed a significant increase in the number of embryos contained within *P. vranovensis*-primed offspring (Figure 1A). To determine whether this accelerated development phenotype could persist for multiple generations, we counted embryos from the F2 offspring of naïve and *P. vranovensis*-infected animals. We observed no difference in the growth rate of these F2 offspring (Figure 1B). Together, these data show that offspring from parents exposed to *P. vranovensis* experience accelerated development, and that these effects last only a single generation.

**Figure 1.**
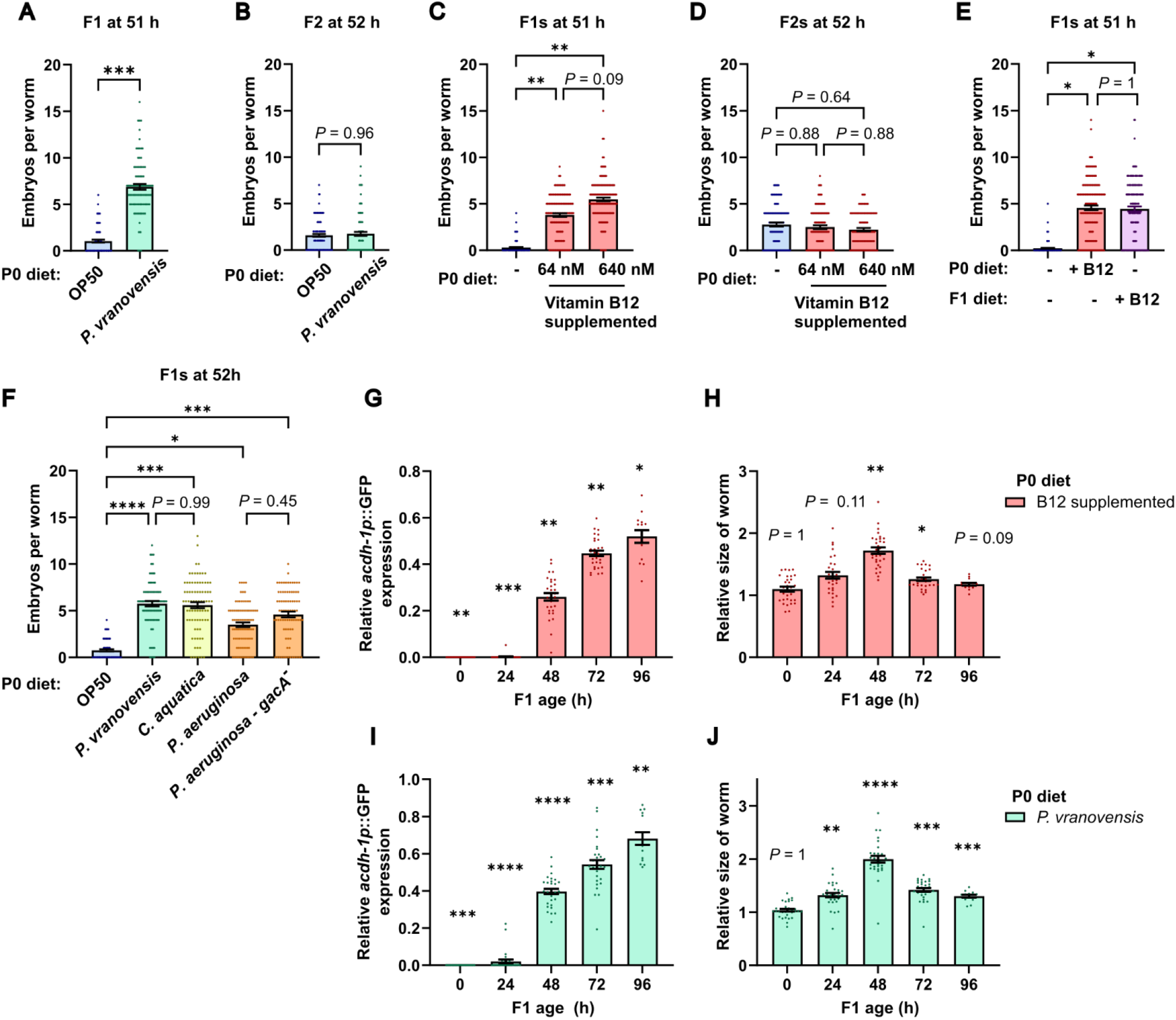
Parental exposure to vitamin B12 induces intergeneration growth acceleration in offspring. (A and B) N2 L4 stage worms were grown on OP50 or *P. vranovensis* for 24 h and their F1 and F2 offspring were grown on OP50. (A) Number of embryos in F1 offspring at 51 h and (B) F2 offspring at 52 h. Data is from three independent replicates of n = 20 to 50 worms. (C-E) N2 L1 stage worms were grown on OP50 or OP50 supplemented with vitamin B12 at a concentration of 64 nM or 640 nM. (C) Number of embryos in F1 offspring at 51 h and (D) F2 offspring at 52 h grown on OP50. Data is from three (C) or two (D) independent replicates of n = 40 to 56 worms. (E) Number of embryos in the F1 offspring grown on OP50 or OP50 supplemented with 640 nM of vitamin B12. Data is from three independent replicates of n = 50 worms. (F) L4 stage animals were grown on different bacterial diets for 24 h and F1 offspring were grown on OP50. Number of embryos in F1 offspring at 51 h. Data is from three independent replicates of n = 20 to 30 worms. (G and I) Levels of *achd-1*p::GFP fluorescence in F1 offspring of worms grown on vitamin B12 (G) or *P. vranovensis* (I), normalized to the offspring of OP50-fed worms at the same timepoint. (H and J) Size of the worms in (G) and (I) normalized to the offspring of OP50-fed worms. Data is from two independent replicates of n = 7 to 15 worms. Horizontal bars represent mean ± SEM. Dots represent individual worms. The *P* values were determined by unpaired two-tailed Student’s t test (A and B), one-way ANOVA with post hoc (C to F), or two-way ANOVA with post hoc (G to J). Significance defined as * p < 0.05, ** p < 0.01, *** p < 0.001, **** p < 0.0001.

Unlike *E. coli* OP50, many *Pseudomonas* bacteria produce vitamin B12. This metabolite can accelerate development in animals directly exposed to it [8]. As such, we next tested whether dietary vitamin B12 may accelerate development in not only the parent, but also the offspring of exposed worms. We plated L4 stage populations of *C. elegans* on either OP50 or OP50 supplemented with 64 nm or 640 nM of vitamin B12 (adenosyl-cobalamin). After 24 h, we isolated embryos and monitored development in the F1 generation by counting embryo production after 51 hours. Offspring from vitamin B12-treated parents had a significant increase in the number of embryos compared to progeny from parents grown on OP50 (Figure 1C). Furthermore, the number of embryos was greater in animals whose parents had been exposed to the higher concentration of vitamin B12. Similar to data obtained for *P. vranovensis*-primed worms, no accelerated development phenotype was observed in the F2 offspring (Figure 1D). To determine if parental exposure to vitamin B12 can provide the same degree of growth acceleration as for animals directly exposed to the metabolite, we compared vitamin B12-primed L1s grown on OP50 to naïve L1s grown on OP50 supplemented with vitamin B12 (Figure 1E). We observed similar embryo numbers from both populations, suggesting that intergenerationally provided vitamin B12 can cause growth acceleration to a similar extent as that provided within the same generation.

We next tested whether several bacterial species known to produce vitamin B12 were able to increase the rate of development in the offspring of exposed parents [8]. Parent populations were either maintained on OP50 or grown on *Comamonas aquatica*, wild-type *Pseudomonas aeruginosa* strain PA14, or *GacA*, an avirulent mutant strain of PA14. F1 offspring from parents grown on any of these other bacterial sources had a significant increase in developmental speed (Figure 1F).

Vitamin B12 has been shown to repress the ‘dietary sensor’ *acdh-1* [9]. To determine whether vitamin B12-primed F1s repress *acdh-1*, we monitored gene expression using the transgenic reporter line *Pacdh-1*::*GFP*. GFP signal was greatly diminished in newly hatched L1s, and *acdh-1* expression was still significantly repressed after 24 h on OP50 (Figure 1G). Though expression of *acdh-1* began to recover after 48 h, *acdh-1* expression in F1s from parents exposed to vitamin B12 was still significantly lower than their naïve counterparts even after 96 h. The F1 offspring of vitamin-B12 exposed worms were significantly larger than their naïve counterparts throughout development and into adulthood (Figure 1H). This increase in size versus naïve F1 worms was most apparent 48 h after hatching. Similar levels *of acdh-1* repression and larger body sizes were observed in the F1 offspring of animals exposed to *P. vranovensis* versus those grown on OP50 (Figures 1I and 1J).

### Dietary vitamin B12 provides tolerance to microsporidia infection

Infection with *N. parisii* causes a decrease in fitness when worms are infected as L1s, but not as L4s, suggesting that faster growth could diminish the negative impacts of infection [14]. To test whether the accelerated development phenotype associated with parental exposure to vitamin B12 provides a fitness advantage to offspring, we infected offspring with *N. parisii*. Here, na**ï**ve, vitamin B12*-*primed, or *P. vranovensis*-primed F1 populations were challenged with a high dose of *N. parisii* at the L1 stage. A population of naïve F1 animals were also grown directly on media supplemented with vitamin B12 and infected with *N. parisii*. Animals were fixed after 72 h and stained with DY96 to visualize both worm embryos and the chitin-containing microsporidia spores. At this timepoint, the number of embryos was slightly (∼20%), but not significantly, elevated in both vitamin B12*-* and *P. vranovensis* -primed F1s under non-infection conditions. However, significant differences in embryo numbers were seen in animals exposed to *N. parisii*, with vitamin B12- and *P. vranovensis-*primed F1s containing four-fold more embryos than naïve worms (Figure 2A). Strikingly, image analysis of stained spores revealed no significant differences in pathogen burden between these F1 populations (Figure 2B). Similar results were observed with naïve F1s grown on vitamin B12 as with offspring that came from parents exposed to vitamin B12 (Figure 2A-B). To determine if the protection to *N. parisii* infection caused by parental B12 exposure increased total reproductive output, we counted all the offspring generated by the F1 animals. We observed no significant difference in the number of offspring produced by naïve and primed worms under non-infection conditions. However, under the stress of microsporidia infection, naïve worms produced on average 6.80 progeny, while vitamin B12- and *P. vranovensis-*primed F1s produced 38.54 and 33.62 progeny, respectively (Figure 2C).

**Figure 2.**
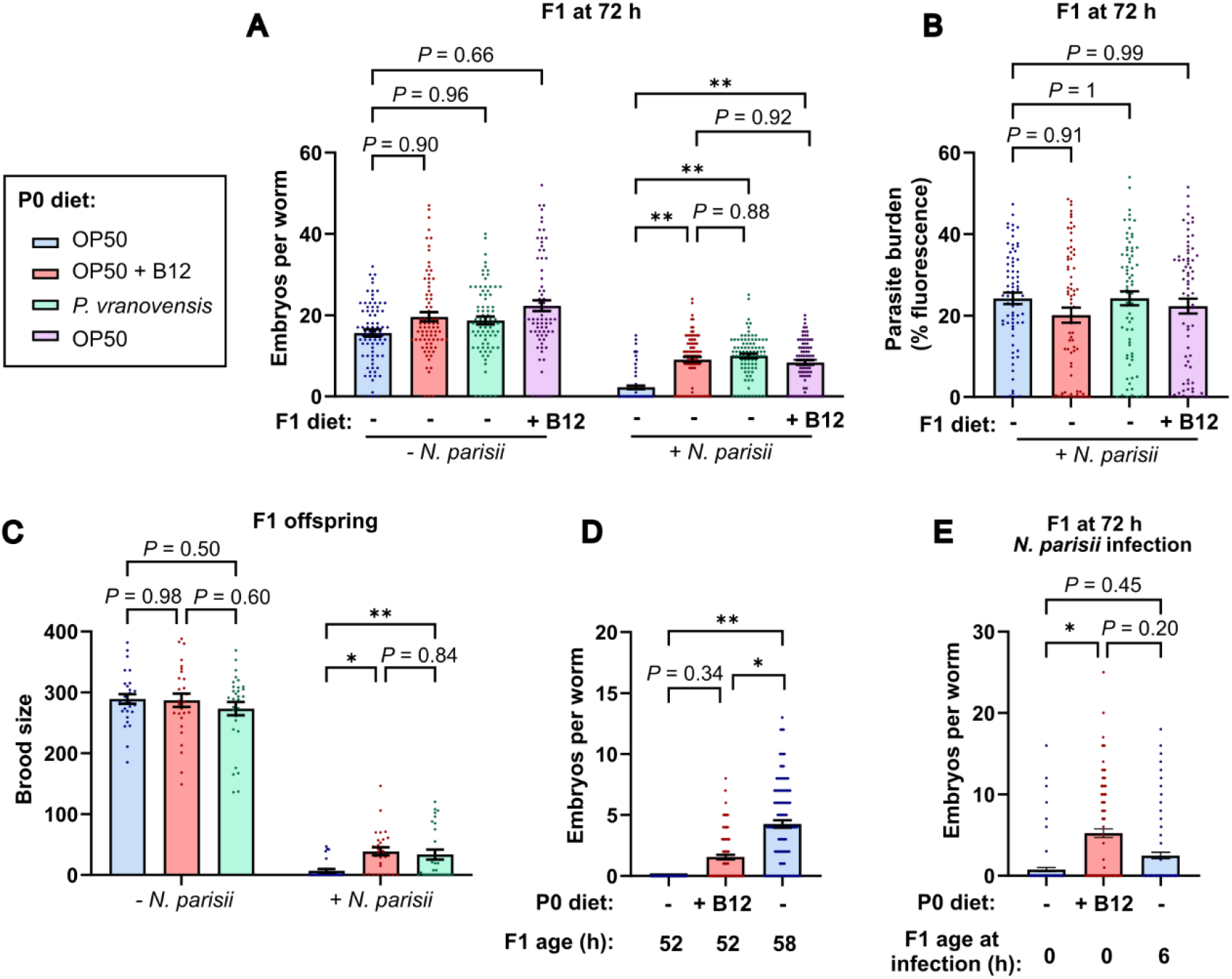
Vitamin B12 provides intergenerational tolerance to *N. parisii* infection. (A-B) L1 stage progeny from parents grown on OP50 or OP50 supplemented with 640 nM of vitamin B12 were either uninfected or infected with *N. parisii* for 72 hours, fixed, and stained with DY96. (A) Number of embryos per worm. (B) Quantitation of DY96 fluorescent spores per worm. Data is from three independent replicates of n = 22 to 30 worms. (C) Brood size of uninfected and infected F1 offspring of worms fed different diets. Data is from three independent replicates of n = 10 worms. (D and E) Number of embryos in naïve and vitamin B12-primed F1 offspring in uninfected (D) and *N. parisii* infection (E) conditions. Data is from four independent replicates of n = 20 to 40 worms. The *P* values were determined by one-way ANOVA with post hoc (A to E). Significance defined as * p < 0.05, ** p < 0.01, *** p < 0.001, **** p < 0.0001.

We next sought to test whether growth acceleration by other means could improve fitness outcomes on *N. parisii*. Naïve worms after 58 h of growth have more embryos than vitamin B12-primed worms at 52 h (Figure 2D). To test if extra time for development could provide similar protection as vitamin B12-priming, we allowed naïve offspring to grow on OP50 for 6 hours prior to initiating *N. parisii* infection. After 72 h, the naïve worms infected at a later developmental age have slightly, but not significantly more embryos than naïve worms not given time to develop before infection. However, infected vitamin B12-primed offspring had more embryos than the older naïve worms (Figure 2E). This suggests that although advanced growth at the L1 stage can result in a small increase in reproductive fitness when infected, the effect of vitamin B12 throughout development is important for tolerance to *N. parisii* infection.

### Intergenerational protection to pathogens provided by vitamin b12 is mediated through multiple *C. elegans* pathways

The effects of vitamin B12 on *C. elegans* occurs through two different pathways [8]. One of these pathways is methionine biosynthesis which is dependent on the methionine synthase METR-1 and is necessary for the growth acceleration caused by vitamin B12. The other vitamin B12-depedant pathway is propionyl-CoA breakdown, which relies on the function of the methylmalonyl-CoA mutase MMCM-1. To test which pathway is responsible for the intergenerational growth acceleration, we grew N2, *mmcm-1*, and *metr-1* animals on either vitamin B12, *P. vranovensis*, or OP50. We tested F1 animals from these parents and observed that vitamin B12-primed or *P. vranovensis*-primed *mmcm-1* mutants displayed increased embryo numbers at 52-54 h, but *metr-1* mutants did not (Figure 3A). In agreement with a role for growth acceleration in protecting against microsporidia infection, when vitamin B12-primed or *P. vranovensis*-primed offspring were infected with *N. parisii, metr-1* mutants did not display an increase in embryo numbers, but *mmcm-1* mutants did (Figure 3B). We observed similar pathogen burden in both mutants and wild-type animals (Figure 3C).

**Figure 3.**
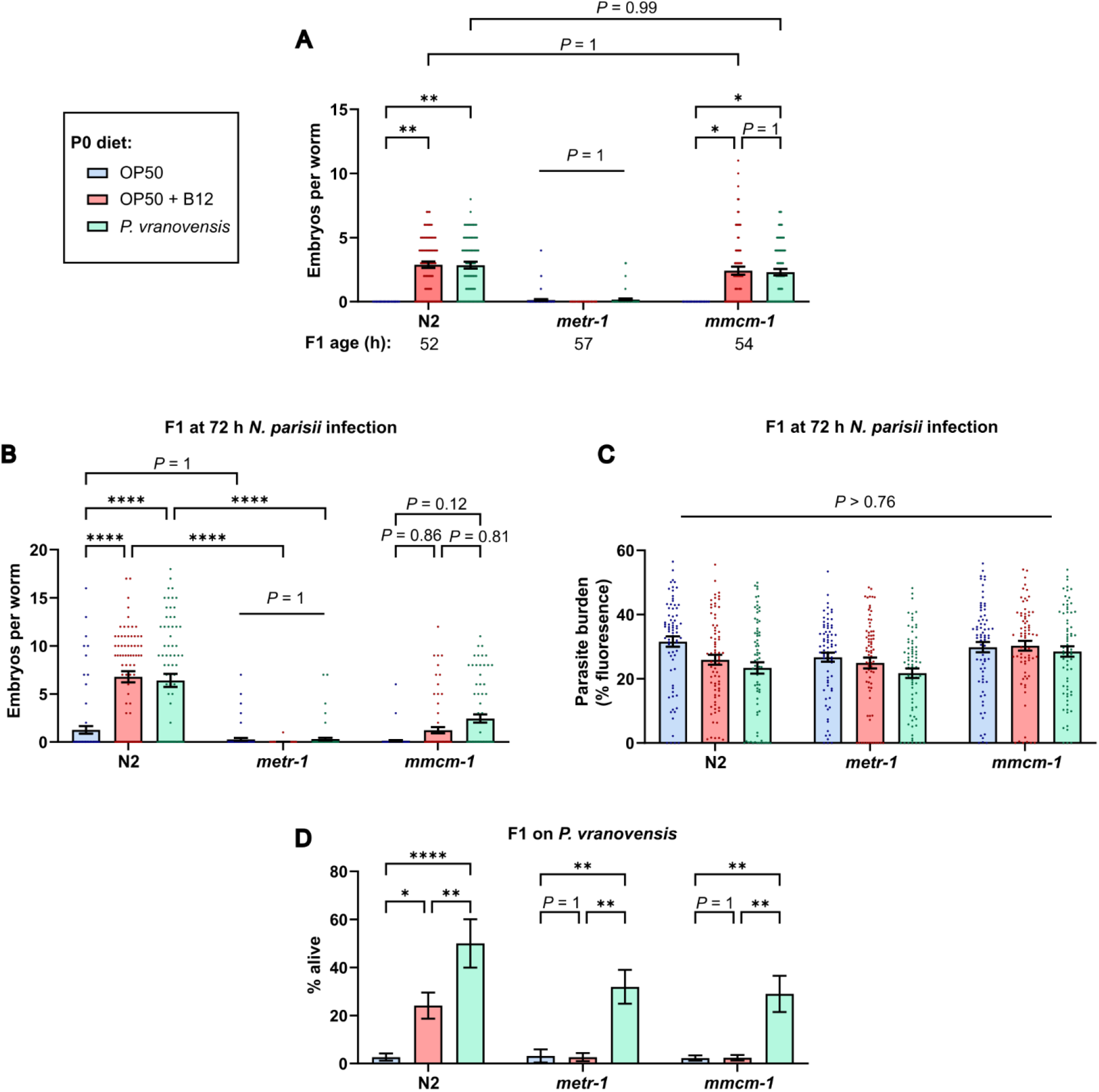
Vitamin B12 pathways provide protection to pathogens. (A to D) N2, *metr-1* and *mmcm-1* P0 worms were fed diets of OP50, OP50 supplemented with vitamin B12, or *P. vranovensis*. (A) Number of embryos in F1 offspring grown on OP50. (B-C) L1 stage offspring were infected with *N. parisii* for 72 h, fixed, and stained with DY96. (B) Number of embryos per worm. (C) Quantitation of DY96 fluorescent spores per worm. Data is from three independent replicates of n = 23 to 25 worms. (D) Offspring embryos were plated on lawns *P. Vranovensis*. Percentage of worms surviving after 24 h. Data is from four independent replicates of n = 100 worms. Horizontal bars represent mean ± SEM. Dots represent individual worms. The *P* values were determined by two-way ANOVA with post hoc. Significance defined as * p < 0.05, ** p < 0.01, *** p < 0.001, **** p < 0.0001.

Vitamin B12 has been shown to protect against killing from *Pseudomonas aeruginosa* through the propionyl-CoA breakdown pathway [15]. Progeny from parents grown on *P. vranovensis* are protected from killing caused by *P. vranovensis*. This adaptation is mediated through upregulation of several genes including, *cysl-1* and *cysl-2*, and animals lacking these genes are no longer intergenerationally protected from killing [4]. To determine if vitamin B12 alone can provide intergenerational protection to *P. vranovensis*, we grew worms on either OP50, vitamin B12, or *P. vranovensis*. We then isolated progeny from these parents and grew the F1s on either OP50 or *P. vranovensis*. Offspring from parents grown on *P. vranovensis* were resistant to killing by *P. vranovensis* and protection to killing was still observed in *metr-1* and *mmcm-1* mutants (Figure 3D). Offspring from parents grown on vitamin B12 were also resistant to killing by *P. vranovensis*. Interestingly, when we infected vitamin B12-primed offspring with *P. vranovensis*, both *metr-1* and *mmcm-1* mutants were not protected from killing (Figure 3D). Together, these results suggest that developmental acceleration from the METR-1 pathway protects animals against *N. parisii*, while both pathways are involved in meditating intergenerational protection against *P. vranovensis*.

### Summary

Here we investigate the intergenerational interplay between pathogens and nutrients. Intergenerational protection against *P. vranovensis* upregulates enzymes that are thought to neutralize pathogen toxins [4,13]. We show that vitamin B12 can provide similar intergenerational protection to bacterial killing through both the propionyl-CoA breakdown pathway as well as the methionine biosynthesis pathway. In contrast to how *N. parisii*-treated parents produce offspring that resist *N. parisii* invasion [5], we show that parents fed vitamin B12 have offspring that exhibit tolerance to *N. parisii* infection. Tolerance, where pathogen burden remains the same but the negative effect on the host is diminished, is increasingly being seen as an important mechanism for limiting the negative effects of pathogens and nutrients such as iron have been show to provide protection against pathogenic bacteria [16]. Our data also show that modest differences in growth speed can result in large differences in reproductive fitness after *N. parisii* infection, suggesting that developmental speed can be a mechanism of tolerance. Together our results show how there can be multiple effects on offspring from parental exposure to microbes as both dietary effects and immune-priming may be happening concurrently (Figure 4). Recently maternal provisioning of sphingolipid was shown to be neuroprotective and together with our results suggests that intergenerational metabolic effects may be common [17,18]. Our results also suggest that potential dietary effects should be taken into account when interpreting how pathogens cause multigenerational immunity [2].

**Figure 4.**
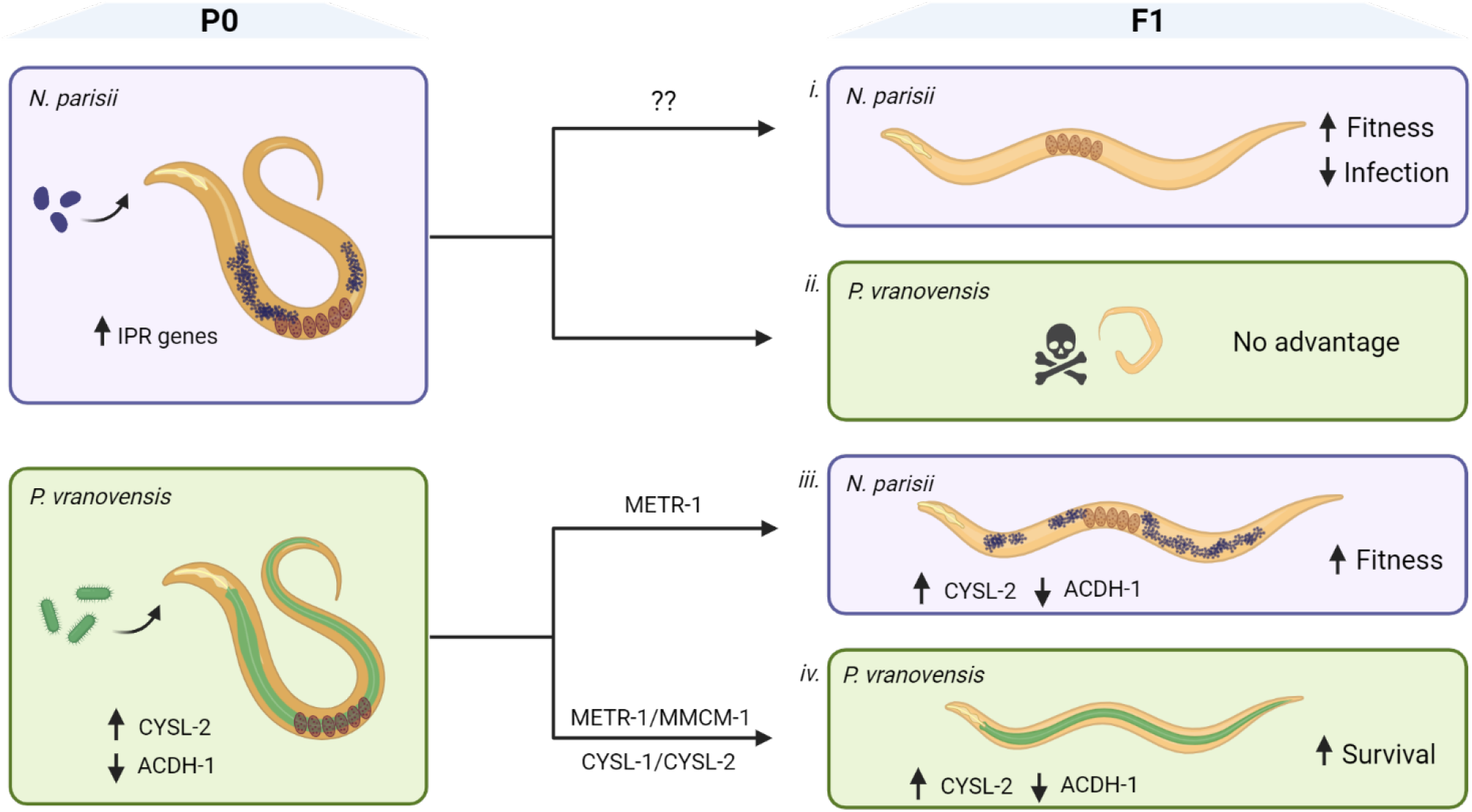
Model of parental pathogen exposure on the outcomes of pathogen-challenged offspring. i. Parental (P0) exposure to *N. parisii* upregulates the intracellular pathogen response (IPR) which is sufficient to produce offspring (F1) that are resistant to *N. parisii* by limiting microsporidia spores in the lumen and preventing invasion [5]. ii. Parental infection with *N. parisii* provides no advantage to offspring exposed to *P. vranovensis* [13]. iii. Growth on bacteria that produce vitamin B12 (such as *P. vranovensis)* results in repression of *acdh-1* which is also repressed in the progeny [9]. Offspring from parents exposed to *P. vranovensis* or vitamin B12 have improved fitness when infected with *N. parisii* and this protection is dependent on METR-1. iv. Parents grown on *P. vranovensis* upregulate cysteine synthases including CYSL-2 and this expression change persists in the offspring [4]. Offspring of parents exposed to *P. vranovensis* or vitamin B12 also display increased survival when exposed to *P. vranovensis*, and this survival is dependent both on CYSl-1/CYSL-2 and METR-1/MMCM-1 [4].

## Acknowledgements

We are grateful to Hala Tamim El Jarkass, Edward James, Yin Chen Wan, and Larry Nguyen for providing helpful comments on the manuscript. We thank Lesley MacNeil for the kind gift of bacterial and *C. elegans* strains. Additional *C. elegans* strains were provided by the *Caenorhabditis* Genetics Center, which is funded by the National Institutes of Health (NIH) Office of Research Infrastructure Programs Grant P40 OD010440. Schematics were created using BioRender.com.

## Data availability

All data is presented in Data S1.

## Methods

### Worm maintenance

Strains of *C. elegans* used in this study are included in Table 1. Worms were grown at 21°C on nematode growth media (NGM) plates that were seeded with 10× OP50-1 *E. coli*, as previously described [5]. OP50-1 was prepared by growing cultures to saturation in lysogeny broth (LB) at 37°C for 16-18 h and concentrating to 10x by centrifugation. To synchronize populations, worms were washed off plates using M9 solution and bleached with sodium hypochlorite/1 M NaOH until the eggs of gravid adults were released into solution. Embryos were then washed 3x in 1 ml of M9, resuspended in 5 ml of M9, and rotated at 21°C for 18-24 h to allow for hatching into L1s. Worms were pelleted by centrifugation in microcentrifuge tubes at 1400*g* for 30 s -1 min.

**Table 1.** *C. elegans* strains used in this study.

| Strain | Genotype | Source |
| --- | --- | --- |
| N2 | Wild-type, Bristol strain | Caenorhabditis Genetics Center (CGC) |
| VL749 | *wwIs24* [*acdh-1p::GFP* + *unc-119(+)*] | Macneil Lab |
| RB755 | *metr-1(ok521)* II | CGC |
| RB1434 | *mmcm-1(ok1637)* III | CGC |

### Culturing of *P. vranovensis, P. aeruginosa*, and *C. aquatica* and preparing lawns for feeding

Overnight cultures of *P. vranovensis, C. aquatica*, and *P. aeruginosa* were prepared by inoculating 3-5 mL of LB with one colony and incubating at 37°C with shaking for 16-18 h. Strains used are listed in Table 2. For diets of *P. vranovensis*, 1 mL of overnight culture was spread on a 10 cm NGM plate and the plate was dried. Plates were prepared on the same day as worms were to be transferred to *P. vranovensis* diets. For diets of *C. aquatica*, 1 mL of overnight culture was spread on a 10 cm NGM plate and incubated at 21°C for 24 h for lawns to develop. *P. aeruginosa* plates were prepared by spreading 125 µL of overnight culture on a 10 cm NGM plate. The plates were incubated at 37°C for 24 hours so lawns can develop, then incubated at 21°C for 8-24 h before worms were plated on them.

**Table 2.** Bacteria strains used in this study.

| Species | Strain |
| --- | --- |
| *E. coli* | OP50-1 |
| *P. vranovensis* | BIGb446 |
| *C. aquatica* | DA1877 |
| *P. aeruginosa* | PA14 |
| *P. aeruginosa* | PA14-*gacA*<sup>-</sup> |

### Priming worms with *P. vranovensis* and other bacteria

To obtain *P. vranovensis*-primed F1s, 2500 bleach-synchronized L1 P0s were mixed with 1 mL of 10X OP50 and plated on 10 cm NGM plates for 48 h incubation at 21°C. The P0s were then washed off the plate and re-plated on an NGM plate seeded with 1 mL of 1X *P. vranovensis* BIGb446 overnight culture. After 24 h incubation at 21°C, the P0s were collected and bleached to obtain *P. vranovensis*-primed F1 embryos. Naïve F1s for comparison were prepared by bleaching P0s that had been incubated with OP50 from L1 for a total of 72 h at 21°C.

To obtain naïve and *P. vranovensis*-primed F2s, 2500 bleach synchronized naïve and *P. vranovensis*-primed F1s were mixed with 1 mL of 10X OP50 and plated on a 10 cm NGM plate. After 72 h incubation at 21°C, the F1s were bleached to release F2 embryos.

Priming with *C. aquatica* and *P. aeruginosa* were done in the same way as described for *P. vranovensis*, with P0 worms transferred to lawns of *C. aquatica* and *P. aeruginosa* after 48 h on OP50.

### Supplementing diets and priming worms with Vitamin B12

Vitamin B12 (Sigma-Aldrich C0884-25MG) at a stock concentration of 633 µM (1mg/mL) was diluted and spread on NGM plates to prepare plates used for vitamin B12-supplemented diets. Unless otherwise specified, the NGM was prepared to have a vitamin B12 concentration of 640 nM. To obtain vitamin B12-primed F1s, 2500 bleach-synchronized P0 L1 worms were mixed with 1 mL of 10X OP50 and plated on NGM with vitamin B12. The plates were dried and incubated at 21°C for 72 h. After incubation, the worms were collected and bleached to obtain F1 embryos. To obtain vitamin B12-primed F2s, 2500 vitamin B12-primed F1 L1s were plated on 10X OP50 for 72 h, then bleached to obtain F2 embryos.

### *acdh-1* reporter assay

Vitamin B12 and *P. vranovensis-*primed *acdh-1*p::GFP worms were generated as described above. Live naïve and primed worms on NGM plates were imaged with a Axio Zoom.V16 (Zeiss) and the amount of GFP fluorescence and size were quantified as described below.

### Growth acceleration assay

To assess changes in growth, 1000 bleach synchronized L1s were mixed with 400 µL of 10X OP50 and plated on 6 cm NGM plates. The plates were dried and incubated at 21°C for 51-52 h for N2 worms or until embryos are visible on the plates for slower growing strains. After incubation, the worms were collected, acetone fixed, and stained with DY96 to visualize the embryos with fluorescence microscopy.

### Preparation of microsporidia spores

Preparation of *N. parisii* (ERTm1) spores was performed as previously described [5]. Briefly, large populations of *C. elegans* N2 worms were infected with microsporidia. Once heavily infected, worms were mechanically disrupted using 1-mm-diameter zirconia beads (BioSpec Products Inc.), and the resulting lysate filtered through 5-μm filters (Millipore Sigma). Contamination checks were performed on spore preparations and those free of contaminating bacteria were stored at -80°C. The concentration of the spores was determined by staining spores with DY96 and using a sperm counting slide (Cell-VU). Each assay was performed using the same batch of spores.

### Microsporidia Infection assays

1000 bleach-synchronized L1s were mixed with a high dose (7.5 million) of *N. parisii* (ERTm1) spores and 400 uL of 10X OP50 and plated on a 6 cm NGM plate. After 72 h incubation at 21°C, worms were fixed in acetone for DY96 staining and analysis.

### Fixation and staining with DY96

Worms were fixed in 1 mL of acetone for 10 min at RT prior to staining. To assay number of worm embryos and determine microsporidia burden, *N. parisii* spores were visualized with the chitin-binding dye DY96. Acetone-fixed animals were washed 2x in 1 ml of PBST, resuspended in 500 μl staining solution [PBST, 0.1% SDS, and DY96 (20 μg/ml)], and rotated at 21°C for 30 min in the dark. For imaging, stained worms were resuspended in 15 μl of EverBrite Mounting Medium (Biotium) and mounted on slides. To pellet worms during staining protocols, animals were centrifuged in microcentrifuge tubes at 10,000*g* for 30 s.

### Microscopy and image analysis

For quantification of embryos, as well as microsporidia burden, worms were imaged using an Axio Imager 2 (Zeiss). Pathogen burden was determined using ImageJ/FIJI [19]. For this, each individual worm was defined as a “region of interest” and fluorescence from DY96-stained microsporidia was subject to “threshold” and “measure area percentage” functions. Threshold values were adjusted to capture the brighter signal from microsporidia spores and exclude the dimmer signal from worm embryos [20]. Final values represent the percentage of an individual worm’s body displaying fluorescence from spores.

For quantification of *acdh-1*p::GFP fluorescence and worm size, worms were imaged on an Axiozoom (Zeiss). On ImageJ/FIJI, individual worms were defined as a “region of interest” and the fluorescence from GFP was subject to the “measure area percentage” function. Worm size was determined from the “measure area” function, which determines the number of pixels contained in the “region of interest”.

### Brood size assay

1000 naïve, *P. vranovensis*-primed, and Vitamin B12-primed F1s were either mixed and plated with 400 µL of 10X OP50 or infected by being mixed and plated with 400 µL of 10X OP50 and 7.5 million ERTm1 spores. At 48 hpi, 10 L4s from each condition were picked individually onto NGM plates seeded with OP50. The worms were transferred to new NGM plates each day for 5 days. F2 offspring from each plate were counted 24 h after the F1 worm was removed from the plate. The brood size is the sum of total F2 offspring from an individual F1 worm.

### Enhanced development assay

To initiate development before infection, 1000 bleach-synchronized L1s hatched for 18 h on rotator were mixed with 6 µL of 10X OP50 and plated on a 6 cm NGM plate. After 6 h of incubation at 21°C, the worms were collected and washed in M9 before infection with *N. parisii*.

### *P. vranovensis* killing assay

As previously described [13], 500 µL of *P. vranovensis* overnight culture was spread onto a 6 cm NGM plate. After drying the plate, ∼100 embryos obtained from bleaching were plated on the *P. vranovensis* lawns. Survival was scored after 24 h at 21°C.

### Statistical analyses

*P*-values were determined by two-tailed unpaired Student’s *t* test or ANOVA. *P*-values not reaching significance are presented in the figures. The calculation of *P*-values was done using Prism software (GraphPad Software Inc.). Statistical significance is defined as *P* < 0.05, unless otherwise stated.

